# The origin and evolution of maize in the American Southwest

**DOI:** 10.1101/013540

**Authors:** Rute R. da Fonseca, Bruce D. Smith, Nathan Wales, Enrico Cappellini, Pontus Skoglund, Matteo Fumagalli, José Alfredo Samaniego, Christian Carøe, María C. Ávila-Arcos, David E. Hufnagel, Thorfinn Sand Korneliussen, Filipe Garrett Vieira, Mattias Jakobsson, Bernardo Arriaza, Eske Willerslev, Rasmus Nielsen, Matthew B. Hufford, Anders Albrechtsen, Jeffrey Ross-Ibarra, M. Thomas P. Gilbert

**Affiliations:** Centre for GeoGenetics, University of Copenhagen, Copenhagen, Denmark.; The Bioinformatics Centre, University of Copenhagen, Copenhagen, Denmark.; Program in Human Ecology and Archaeobiology, Department of Anthropology, National Museum of Natural History, Smithsonian Institution, Washington D.C., USA.; Department of Genetics, Harvard Medical School, Boston, USA.; Department of Integrative Biology and Statistics, University of California, Berkeley, USA.; Department of Evolutionary Biology, Uppsala University, Uppsala, Sweden.; Science for Life Laboratory, Uppsala University, Uppsala, Sweden.; Instituto de Alta Investigación, Universidad de Tarapacá, Arica, Chile.; Department of Ecology, Evolution, & Organismal Biology, Iowa State University, USA.; Department of Plant Sciences, Center for Population Biology and Genome Center, University of California, Davis, USA.; Trace and Environmental DNA Laboratory, Department of Environment and Agriculture, Curtin University, Perth, Australia.

## Abstract

Maize offers an ideal system through which to demonstrate the potential of ancient population genomic techniques for reconstructing the evolution and spread of domesticates. The diffusion of maize from Mexico into the North American Southwest (SW) remains contentious with the available evidence being restricted to morphological studies of ancient maize plant material. We captured 1 Mb of nuclear DNA from 32 archaeological maize samples spanning 6000 years and compared them with modern landraces including those from the Mexican West coast and highlands. We found that the initial diffusion of domesticated maize into the SW is likely to have occurred through a highland route. However, by 2000 years ago a Pacific coastal corridor was also being used. Furthermore, we could distinguish between genes that were selected for early during domestication (such as *zagl1* involved in shattering) from genes that changed in the SW context (e.g. related to sugar content and adaptation to drought) likely as a response to the local arid environment and new cultural uses of maize.

---

Maize was domesticated from the wild grass teosinte in southern Mexico more than 6000 years ago ^1–3^. One of the key stepping stones in maize’s spread and evolution was its entry into the area today encompassed by the United States (USA) via the SW around 4100 years before present ^4^. In the SW, maize adapted to the local arid environment and to new cultural uses ^4^, with changes reflected in evolving cob morphology. We have used genetic information from archaeological SW maize, together with that of ancient maize samples from Chile and Mexico (including the third oldest maize macrofossil ever dated, ca. cal. 5910 BP, from Tehuacan), to investigate the route of maize diffusion into the SW, and to identify genes presenting evidence of strong directional selection by farming societies of this region.

Using a hybridization-target capture approach ^5^ we generated sequence data spanning predominantly the exons of 348 genes (selection criteria in ^6^; Supplementary Tables 8 and 9) in i) 3 ancient Mexican samples dating to ca. cal. 5910 BP, 5280 BP and 1410 BP, ii) 25 temporally distributed ancient SW maize samples dating to ca. cal. 4000-3000, 2000 and 750 BP (SW3K, SW2K and SW750 hereafter), and iii) 4 ancient Chilean samples dating to less than 1500 BP (Table 1). Additional sequence data from a previous study ^5^ was added to the dataset (one ancient sample from Mexico, Supplementary Table 7).

**Table 1.** Identification and summary statistics of the ancient samples sequenced in this study.

| Age group | Type of analyses* | Ids | Intercept of radiocarbon age with calibration curve years BP | Cob morphology (P: pineapple; C:cylinder) | Site | Retained nucleotides | Average depth (targets) |
| --- | --- | --- | --- | --- | --- | --- | --- |
| SW3K | a,d | SW443 | 2,780 |  | McEuen, USA | 8,230,593 | 13 |
|  | t,a,d | SW4Ba | 3,390 |  | Bat Cave, USA | 17,621,611 | 9 |
| SW2K | a,d,s | SW207 | 1,860 | P, 12 row, 2.0 cm | Tularosa, USA | 5,870,362 | 11 |
|  | s | SW256 | ca. 1,850-1,750 | P, 12 row, 2.5 cm |  | 3,768,546 | 9 |
|  | s | SW261 | ca. 1,850-1,750 | P, 10 row, 1.9 cm |  | 4,613,152 | 9 |
|  | a,d,s | SW264 | 1,820 | P, 12 row 1.8 cm |  | 15,134,398 | 14 |
|  | s | SW278 | ca. 1,850-1,750 | P, 12 row, 2.2 cm |  | 3,431,137 | 10 |
|  | a,d,s | SW280 | ca. 1,850-1,750 | P, 10 row, 2.1 cm |  | 5,209,183 | 6 |
|  | s | SW283 | 1,860; 1,850; 1,830 | P, 12 row, 2.2 cm |  | 5,642,954 | 3 |
|  | s | SW288 | ca. 1,850-1,750 | P, 12 row, 2.3 cm |  | 148,791 | 1 |
|  | s | SW296 | ca. 1,850-1,750 | P, 10 row, 1.9 cm |  | 2,072,254 | 5 |
|  | t,a,d,s | SW298 | 1,770; 1,760; 1,740 | P, 12 row, 2.0 cm |  | 80,568,726 | 10 |
| SW750 | s | SW105 | 670 | C, 10 row, 1.5 cm | Tularosa, USA | 2,058,626 | 4 |
|  | a,d,s | SW107 | ca. 700-900 | C, 8 row, 1.3cm |  | 34,929,483 | 20 |
|  | a,d,s | SW109 | ca. 700-900 | C, 8 row, 1.3 cm |  | 12,364,145 | 15 |
|  | a,d,s | SW110 | ca. 700-900 | C, 8 row, 1.6 cm |  | 35,088,565 | 17 |
|  | a,d,s | SW111 | ca. 700-900 | C, 8 row, 1.5 cm |  | 29,640,515 | 19 |
|  | a,d,s | SW112 | ca. 700-900 | C, 8 row, 1.4 cm |  | 22,887,209 | 16 |
|  | s | SW118 | ca. 700-900 | C, 8 row, 1.2 cm |  | 3,855,808 | 4 |
|  | a,d,s | SW121 | 790 | C, 10 row, 1.5 cm |  | 29,736,402 | 7 |
|  | a,d,s | SW124 | ca. 700-900 | C, 8 row, 1.3 cm |  | 33,518,448 | 18 |
|  | a,d,s | SW132 | 740 | C, 8 row, 1.4 cm |  | 17,131,288 | 18 |
|  | t,a,d,s | SW146 | 690 | C, 8 row, 1.3 cm |  | 111,329,149 | 12 |
|  | s | SW1b9 | 740 | C, 8 row, 1.5 cm |  | 686,34 | 2 |
|  | a | SW1AX |  |  | Turkey House, USA | 59,526,622 | 25 |
|  | a | TH563 | 5,910 | 4 Ranks, 8 Rows, 1.2 cm | Tehuacan, Mexico | 9,544,881 | 3 |
|  | t,a | TH564 | 5,280; 5,160; 5,140; 5,100 | 4 Ranks, 8 Rows, 1.1 cm |  | 10,791,297 | 5 |
|  | a | TH157 | 1,410 | 8 Rows, 1.5 cm |  | 18,126,654 | 2 |
|  | a | AR14B |  |  | Arica, Chile | 5,328,366 | 16 |
|  | a | AR1A9 |  |  |  | 11,261,584 | 11 |
|  | a | AR1A8 |  |  |  | 286,639,854 | 24 |
|  | a | AR171 |  |  |  | 159,400,189 | 21 |

\*t: Treemix (Fig. 1A), a: NGSadmix (Fig. 1b, Supplementary Fig. 5), d: D-statistics (Fig. 1c, Supplementary Fig. 12), s: selection tests (Fig. 2, Fig. 3, Supplementary Fig. 10)

The sequence data display typical ancient DNA (aDNA) damage patterns (Figs. S1 and S2). Error rates for the SW750 and SW2K individuals were below 0.2% per base after removal of C to T and G to A transitions (potentially resulting from aDNA damage; Supplementary Fig. 3). Comparisons were made with sequence data from modern maize landraces and the wild teosintes *Zea mays* ssp. *parviglumis* (henceforth parviglumis, the grass from which maize was domesticated) and *Z. m*. ssp. *mexicana* (henceforth mexicana) from maize HapMap2 ^7^ and shotgun data from an open-pollinated Mexican highland individual ^6^ (Supplementary Fig. 4 and Supplementary Tables 4 and 5).

**Figure 1.**
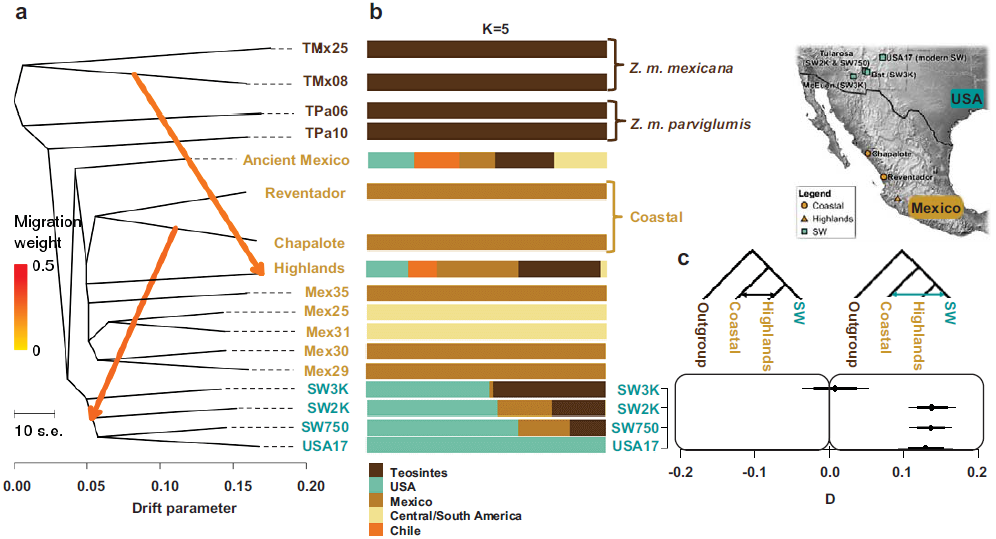
Origins of the SW ancient maize samples. SW3K, SW2K and SW750 correspond to SW maize from ∼3000, ∼2000 and ∼750 BP. The ancient Mexican sample dates to 4,500 BP (TH564). The Mex prefix indicates modern Mexican samples that also include the Reventador, Chapalote and Highland landraces. More details in Table 1 and Supplementary Tables 4 and 5. **a)** Treemix maximum likelihood tree depicting the expected signal of gene flow from *Z. m. Mexicana* into the highland landraces (also Supplementary Fig. 12) and gene flow from the coastal Chapalote into the SW2K. **b)** Subset of the population structure plot determined by NGSadmix with K=5 ^6^ (full plot in Supplementary Fig. 5); each individual is represented by a stacked column of the five proportions. **c)** The results from allele frequency-based *D*-tests^20^ suggest an initial highland diffusion route from Mexico to the SW of the USA followed by extensive gene flow from the Pacific coast Chapalote race (Supplementary Table 6, Supplementary Fig. 12); positive values of D indicate gene flow from the coastal varieties into the SW maize; thick and thin bars correspond to 2 and 3 standard errors, respectively.

**Figure 2.**
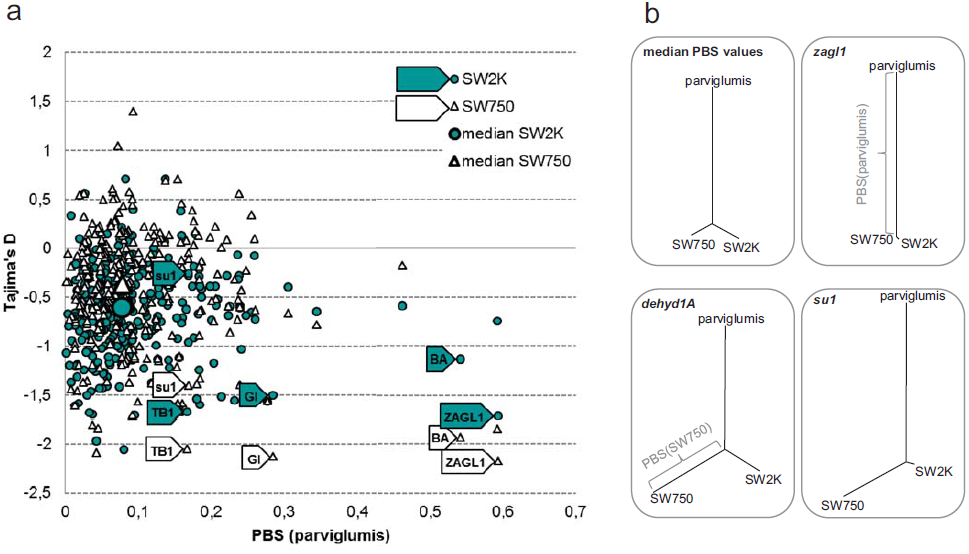
Potential targets of selection during domestication. **a)** Tajima’s D for the two SW populations dated to ∼2000 (colored dots) and ∼750 BP (white triangles) plotted against the PBS distance for parviglumis. *zagl1* shows the highest dissimilarity between parviglumis and the ancient Southwest landraces, i.e. the largest PBS(parviglumis). The gene with the lowest Tajima’s D value for the SW750 population is also *zagl1*. Genes with major roles in domestication traits are depicted in trapezoids. **b)** Gene trees built using PBS distances. *dehyd1A* is the top outlier for PBS(SW750) (Supplementary Fig. 10) and *su1* displayed the highest decrease in nucleotide diversity between the SW2K and the SW750 populations.

**Figure 3.**
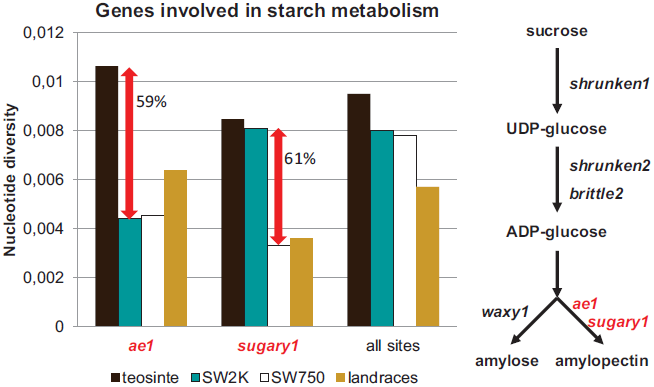
Timing of selective pressures on genes involved in the starch metabolism. Nucleotide diversity variation for two key elements of the starch metabolism pathway. There is a steep decrease in nucleotide diversity before 2,000 BP for *ae1*, whereas the reduction in π for *sugary1* to less than half occurred after 2K and before 750 BP.

The origin of maize in the SW has long remained contentious. For many years the only available archaeological samples of maize were from higher altitude sites, suggesting a high elevation Sierra Madre Occidental corridor of diffusion ^8,9^. More recently discovered samples from lower elevation river valleys ^10,11^ and the morphological similarities between archaeological samples and extant Mexican landraces ^12,13^, however, suggest diffusion along a low-elevation Pacific coastal route. Results from four different analytical approaches indicate that while maize originally entered the SW through a highland route, it subsequently received genetic contributions arriving via the Pacific coastal corridor by 2000 BP: i) Treemix results using ancient and extant samples show gene flow from the Pacific coast Chapalote landrace into the SW2K maize (Fig. 1a); ii) admixture analysis ^6,14^ of ancient and extant samples for k=5 groups shows evidence for an increase in admixture with coastal lowland maize by 2000 BP (Fig.1B, Supplementary Fig. 5), iii) explicit comparison of the different chronological SW groups using the D-statistics ^15^ with ancient and extant samples provides strong support for significantly more shared derived variants between SW individuals and the Pacific coast Mexican lowland race Chapalote than between those SW individuals and the highland landrace Palomero de Jalisco, except for the SW3K group (Fig. 1c), and iv) additional analyses using a panel of 2310 modern maize landraces and teosinte shows that all of SW maize landraces (across elevations) share ancestry with landraces from highland Mexico, highland Guatemala and, to a lesser extent, with mexicana (Supplementary Fig. 6). The admixture analysis (Fig. 1b, Supplementary Fig. 5) also shows evidence of teosinte admixture in all ancient SW maize. Teosinte has no history of occurrence in the SW and was likely absent during the timeframe of the introduction of maize ^16^. Instead, admixture with teosinte almost certainly predated the introduction of maize into the SW, and is evident in the oldest Mexican samples sequenced (Fig. 1b, Supplementary Fig. 5). Furthermore, we interpret the lack of observed admixture with teosinte or Mexican maize in the extant SW Santo Domingo landrace (USA17) to be a result of recent extensive gene flow with other American landraces (Supplementary Fig. 5).

The SW2K and SW750 sample sets present several features that allowed us to identify the targets and timing of prehistoric selection: i) the error rate for these individuals was below 0.2% per base, ii) they correspond to distinct occupations of the same cave site with sample sizes ranging from 10 (SW2K) to 12 (SW750) individuals, which allowed us to calculate various population summary statistics such as F_ST_ and Tajima’s D to time the onset of early genetic changes ^17,18^, iii) the median F_ST_ across all captured genes was 3%, indicating low differentiation. We used the population branch statistic, PBS ^19^, to detect genes that were likely early targets of maize domestication, potentially before arriving in the SW, as they present the highest dissimilarity between parviglumis (HapMap2 individuals, Supplementary Table 4) and ancient SW landraces (Fig. 2a). Seven of the top 10 genes (Supplementary Table 1) are located in regions that were detected as being under selection in an unrelated study by Hufford *et al* ^20^ (the function of the other 3 genes is not well characterized). The top gene, *zagl1*, corresponds to a MADS-box transcription factor associated with shattering, a key domestication feature strongly selected for by human harvesting ^21–23^. The other genes are also well known from their roles in domestication traits: i) *ba1* has a major role in the architecture of maize ^24^, ii) *zcn1* and *ZmGI* are associated with the regulation of flowering ^23,25^, and iii) *tga1* controls the change from encased to exposed kernels ^26^. To provide further evidence of selection we calculated Tajima’s D for each gene ^27^. The median D for the SW750 population is more positive than that of the SW2K population (Supplementary Fig. 8 and Supplementary Table 2), consistent with reduced estimates of effective population size (Supplementary Fig. 9), but many domestication-related genes show the opposite trend, suggesting the influence of recent strong selection (Fig. 2a).

We have also identified a number of candidate loci that appear to have been targeted by selection starting sometime between 2000 and 750 years BP in the context of the SW. The top PBS outlier for SW750 is a dehydration-responsive element-binding protein shown to be up-regulated as much as 50-fold in maize roots under drought conditions ^28^, perhaps a signature of adaptation to arid SW conditions (Supplementary Fig. 10). Furthermore, *ae1* shows reduced diversity in both the SW2K and SW750 populations, consistent with early selection (Fig. 3). Mutations in both *sugary1* locus (*su1*) and *ae1* (Fig. 3) affect the structure of amylopectin ^29^, which is involved in the pasting properties of maize tortillas and porridge ^30^.

While analysis of individual ancient samples suggests the presence of modern maize haplotypes at the *su1* as early as 4400 BP ^17^, our data show a substantial (61%) reduction of diversity at this locus only after 2000 BP (Fig. 3), a signature that correlates with a shift toward larger cobs and floury kernel endosperm ^11^ in archaeological maize around AD 800-1000. The *su1* mutation with the highest allele frequency difference between SW and modern individuals is known to cause the partial replacement of starch by sugar in sweet corn ^31^. Several Native American tribes grew sweet corn before the arrival of Europeans and the high frequency of a *su1* mutation in SW maize could help explain the early appearance and maintenance of sweet corn varieties by Native Americans.

The study of domestication and early crop evolution has largely been limited to the identification of key phenotypic and morphological changes between extant crops and their wild relatives. As demonstrated here, the application of new paleogenomic approaches to well-documented temporal sequences of archaeological assemblages opens a new chapter in the study of domestication: it is now possible to move beyond a simple distinction of "wild" vs. "domesticated" ^32^ and track sequence changes in a wide range of genes over the course of thousands of years.

## Acknowledgments

The authors acknowledge the following grants: Marie Curie Actions IEF 272927, Danish National Research Foundation DNRF94, Danish Council for Independent Research 10-081390 and 1325-00136,Lundbeck Foundation grant R52-A5062, Vand Fondecyt Grant 1130261, a grant from the UC Davis Genome Center for the highland maize sequence. RF is supported by a Young Investigator grant (VKR023446) from Villum Fonden. PS was funded by the Wenner-Gren foundation. The authors thank Ângela Ribeiro, Shohei Takuno and Philip Johnson for comments and discussion and the Danish National High-Throughput DNA Sequencing for technical support.

